# Optogenetic single-cell control of transcription achieves mRNA tunability and reduced variability

**DOI:** 10.1101/142893

**Authors:** Marc Rullan, Dirk Benzinger, Gregor W. Schmidt, Ankit Gupta, Andreas Milias-Argeitis, Mustafa Khammash

## Abstract

The study of gene expression at the single-cell level has exposed the importance of stochasticity for the behavior of cellular systems. Research on cellular variability has mostly relied on observing expression either in response to natural stimuli or to constant gene regulators. However, the ability to probe cells individually can lead to a deeper understanding of the underlying process. Here, we propose an experimental platform for optogenetic feedback control of individual cells. It consists of a digital micromirror device that, coupled to a microscope, can target light-responsive cells with individualized illumination profiles, thereby exploiting the good spatial resolution of optogenetic induction. Together with an automated software pipeline for segmentation, quantification and tracking of single cells, the platform enables independent and real-time control of numerous cells. We demonstrate our platform by regulating transcription in over a hundred yeast cells simultaneously, while achieving tunability of mRNA abundance. Using a novel technique to measure extrinsic variation, we further show that single cell feedback regulation of this highly stochastic process achieves a 10-fold reduction of extrinsic variation in nascent mRNA over population control, with superior control loop properties. Our platform establishes a new, flexible method for studying transcriptional dynamics in single cells.

## Introduction

The study of stochastic gene expression at the single-cell level transformed the field of genetics by revealing and explaining a host of cellular properties previously inaccessible by bulk analysis of cell populations^1–3^. This benefitted from a strong interplay between theory and experiment, with the mathematical analysis of gene expression variation^4,5^ driving biological discovery and vice versa^6–8^.

The vast majority of single-cell gene expression studies have been carried out under uniform induction conditions. Application of the same chemical input to populations of isogenic cells has enabled the quantification of intrinsic and extrinsic components in gene expression variability^8,9^, and provided important insights into the molecular origins of cell-to-cell variations^10^. However, the quantitative study of the different gene expression steps, as well as the processing and propagation of information in gene networks would greatly benefit from probing single cells *individually*^11^. While technically very difficult to attain with chemical inducers at high throughput, induction of single cells with individual dynamic input signals can be effectively achieved by the combination of optogenetics, live single-cell microscopy and patterned illumination systems^12–15^.

Such an experimental platform allows accurate spatial and temporal targeting of single cells with light inputs to study cell-to-cell variability in great detail. Furthermore, real-time cell tracking and quantification of key variables of interest from microscopy images enables the implementation of *in silico* feedback control strategies^16–18^. In such strategies, real-time single-cell measurements are fed into a computer-implemented control algorithm that computes an individual light input for each cell in order to achieve a pre-specified tracking objective. Feedback regulation of single cells opens up possibilities for studying stochastic gene expression dynamics and the consequent cell-to-cell variability at a previously unattainable level of detail.

Previous attempts at single-cell feedback control have been scarce, and of low-throughput. In ref. 19, light-gated recruitment of proteins on the membrane of single mammalian cells was regulated with feedback control. Those experiments were carried out in a timescale of a few seconds, with continuous observation and actuation of a single cell at a time. On the other hand, ref. 18 presented long-term feedback regulation of Hog1-responsive gene expression in single yeast cells by modulating the osmolarity of the cellular environment inside a microfluidic chip. However, since the input signal was applied to all cells simultaneously, only one cell could be tracked and controlled during each 15-h long experiment.

Overcoming the limitations of previous studies, we present here a fully automated experimental platform for fast real-time, long-term individual control of transcription in ~100 yeast cells harboring an optogenetic gene expression system. Transcriptional output of single cells is quantified by an RNA detection system based on the binding of fluorescently-tagged PP7 phage coat protein to RNA stem-loops^20^. In this way, active transcription of a gene of interest can be visualized as a fluorescent nuclear spot whose instantaneous intensity reflects the current nascent RNA load. To achieve single-cell stimulation, a Digital Micromirror Device (DMD) is used to project arbitrary light patterns through the microscope and onto a microfluidic-grown yeast culture^12^. Single-cell light inputs are in turn computed with a software pipeline that acquires and processes microscopy images to track cells and quantify nascent mRNA counts with a time resolution of two minutes.

We use our setup to study the differences between single-cell and population-level feedback control by regulating the nascent mRNA counts of individual cells at pre-specified levels over several hours. We further introduce a novel method based on time-course measurements for the quantification of extrinsic and intrinsic sources of cell-to-cell variability, with which we demonstrate that single-cell control effectively cancels output variation generated by slowly-varying extrinsic fluctuations. This dramatic reduction of extrinsic variation in nascent mRNA counts across the population in turn gives rise to tighter mRNA distributions. The ability to precisely direct the behavior of single-cells, together with the rich datasets generated by our platform form a powerful combination that opens up new possibilities for both the experimental and theoretical study of gene expression in single cells.

## Results

### An experimental setup for single-cell optogenetics

We have built an experimental platform tailored for spatially independent photoinduction of gene expression or signaling in hundreds of single yeast cells in parallel (Figure 1A). To address cells with light, we made use of a projector based on a Digital Micromirror Device^12^ (Methods). The DMD contains an array of about a million individual micromirrors, with each mirror being independently switchable between an “on” and an “off” position. When “on”, the mirror reflects the light of an LED source onto the specimen, while intermediate light intensities can be achieved by fast pulse-width modulation of the mirror position. Coupled with a microscope at sufficient magnification (Supplementary Note 1), the high pixel density of the DMD-based projector can thus achieve a micrometer spatial resolution. This in turn enables the generation of light patterns that can precisely target individual yeast cells within a tightly-packed micro-colony with inputs of arbitrary duration and intensity (Figure 1B).

**Figure 1:**
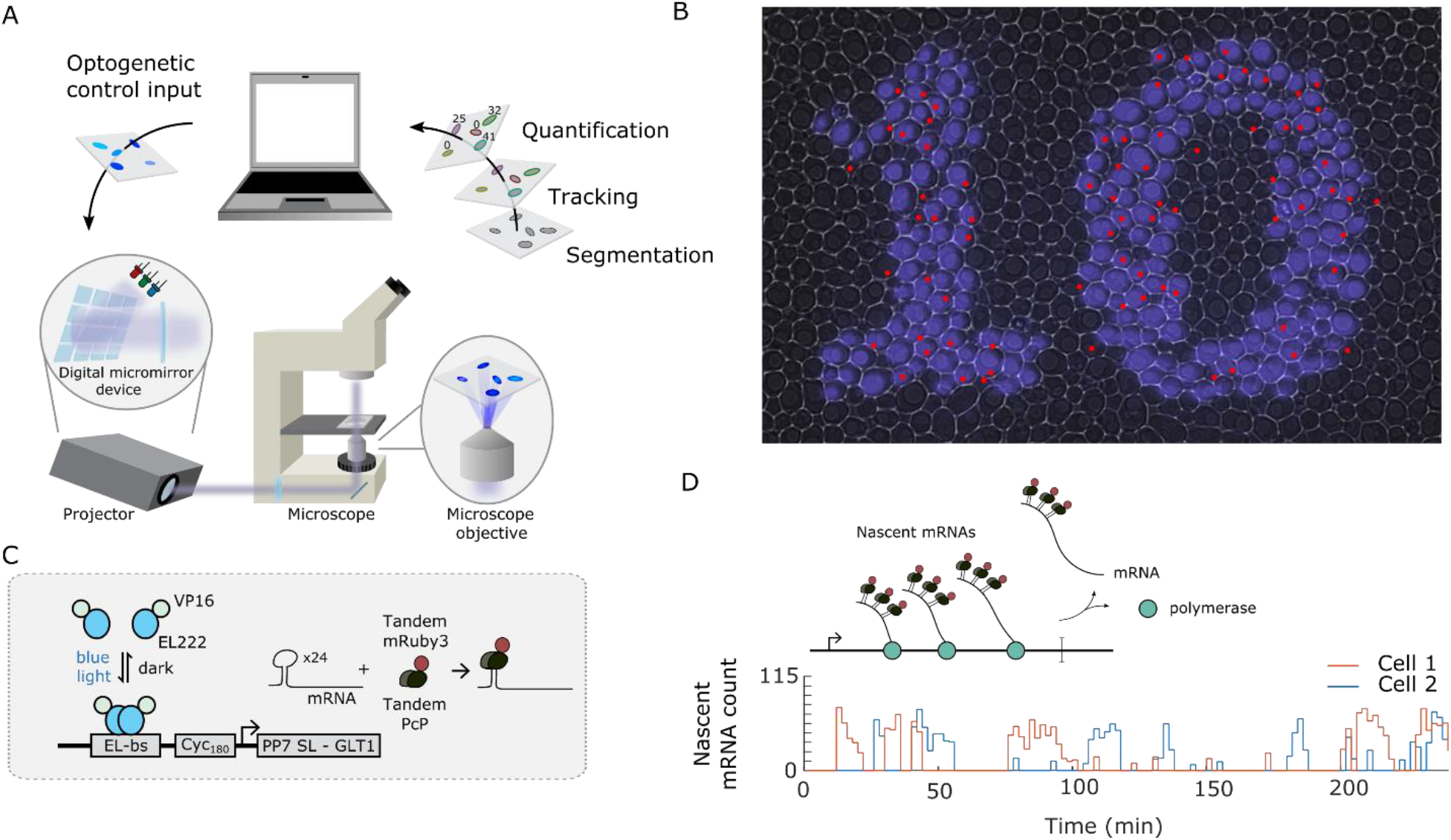
Experimental setup for optogenetic feedback control of single cells. *A: Experimental feedback loop for optogenetic single-cell control. Light-responsive cells are grown under a microscope and periodically imaged to measure the controlled variable of interest. The images are read by a computer in charge of cell segmentation and tracking, and quantification of the cellular readouts. The results are provided to individual feedback controllers (each assigned to a single cell), which compute the light intensity to be projected onto each cell at the next time-point, in order to attain a pre-specified level of the controlled output. The calculated inputs are passed to a digital micromirror device-based projector, responsible for precisely targeting light onto the cells. B: Optogenetic induction of transcription in single-cells. Yeast cells densely growing in a monolayer are illuminated through the DMD-based projector (blue) in the pattern of a number “10”. The active transcription site of each cell (imaged in the fluorescence channel) is marked by a red spot (Supplementary Note 1). Given that transcription of this gene takes place in bursts (see below), it is expected that not all cells exposed to light are actively producing the measured mRNA. C: Optogenetic induction of transcription and mRNA labelling. EL222 homodimerizes in presence of blue light, exposing its DNA-binding domain. The dimer then binds to its cognate promoter, a fusion of five EL222 binding sites (EL-bs) to a 180 base-pair long sequence of the CYC1 promoter (CYC_180_), promoting the expression of a downstream gene due to the activation domain VP16. The regulated gene encodes the GLT1 mRNA with twenty four PP7 stem-loops (PP7 SL) inserted at its 5’ end. These stem-loops are recognized and bound by a tandem dimer of the PP7 bacteriophage coat protein (tdPCP). The fusion of two copies of mRuby3 to tdPCP enables the visualization of the produced mRNAs in live cells. D: Nascent mRNA quantification and experimental demonstration of transcriptional bursting. (Upper) The accumulation of multiple fluorescently-labeled mRNAs actively transcribed (nascent mRNAs) at the transcription site generates a diffraction-limited fluorescent nuclear spot clearly visible under the microscope. The intensity of this spot is proportional to the number of nascent mRNAs present at the transcription site at the time of imaging. (Lower) Nascent mRNA profile in two cells exposed to non-saturating, constant blue light. The cellular response to the stimulus clearly shows that transcription takes place in bursts. Interestingly, the light intensity projected onto the cells does not affect the magnitude of the transcriptional bursts, but rather modulates the fraction of cells transcriptionally active (Supplementary Note 2*).

The parallel, single-cell regulation of transcription across a fast-growing cellular population poses challenges with respect to cell segmentation and tracking, which we overcame by constructing a software pipeline for imaging automation, real-time image processing and light input application (Methods). With this setup, pre-specified temporal and spatial light patterns can be applied to individually tracked cells or cell groups (*open-loop control*). Furthermore, monitoring transcriptional activity within each cell with an mRNA detection system (see below) allows the calculation of light inputs based on the current and past measurements from each cell, in order to achieve a target activity level (*feedback control*). This further required the addition of computational algorithms to quantify the cellular readouts and control algorithms to compute the necessary light input adjustments within our software pipeline (Methods).

Thanks to the careful optimization of all hardware and software components within our feedback loop, our system is capable of updating the light inputs to ~100 tracked yeast cells every two minutes – a frequency that allows real-time feedback control of fast cellular processes such as transcription or signaling.

### Optogenetic control of transcription and nascent mRNA quantification

To regulate transcription using light stimulation we engineered S. *cerevisiae* to include a light-switchable transcription factor (VP-EL222), composed of the LOV-domain protein EL222^21^ fused to the VP16 activation domain (Methods). Blue light stimulation induces conformational changes in EL222, allowing it to homodimerize, bind to its cognate promoter sequence and initiate the transcription of a target gene downstream of a VP-EL222 responsive promoter (Figure 1C).

To obtain a fast readout of the fluctuations in transcriptional activity, we used the PP7-based RNA detection system^20^ to image single-cell transcription events in real time. The system consists of an array of 24 stem-loop encoding sequences fused in front of the target gene (GLT1, Figure 1C). Each stem-loop is recognized and bound by a tandem dimer of the PP7 bacteriophage coat protein fused to the mRuby3 fluorescent protein. As nascent glt1 RNAs get bound by multiple PP7 fusion copies, active transcription sites can be detected as diffraction-limited nuclear fluorescent spots (Figure 1B,D). The number of nascent mRNAs being actively transcribed at a given time can thus be estimated by quantifying the brightness of these spots, after the appropriate calibration^7^ (Methods).

### From population to single-cell control

Applications of feedback control in biology commonly focus on controlling the average behavior of a group of cells^22,23^, which we will refer to as *population control.* In this case, the cells are either measured in bulk, or the single-cell measurements are averaged and fed to a common controller, which in turn determines a common input profile to be applied to all cells (Figure 2A). Since population control is only able to produce one input stimulus seen by all cells, it cannot shape the response of individual cells. While this may steer the average behavior to a desired level, the highly variable dynamic behavior of single cells^24^ means that a large number of cells will have a response that is very different from the average.

In contrast, in the setting of *single-cell control,* each cell is steered independently via a separate controller that is updated only with measurements from that cell (Figure 2A). Here the control objective may be common for all cells or it could vary from cell to cell. This strategy requires measurement, tracking and independent stimulation for each cell. The increased complexity of this task in comparison to population control comes with a benefit: when all controlled systems are not identical, as in the case of living cells^8,24^, application of independent inputs to different cells can compensate for their differences and achieve a much more homogeneous population response.

**Figure 2:**
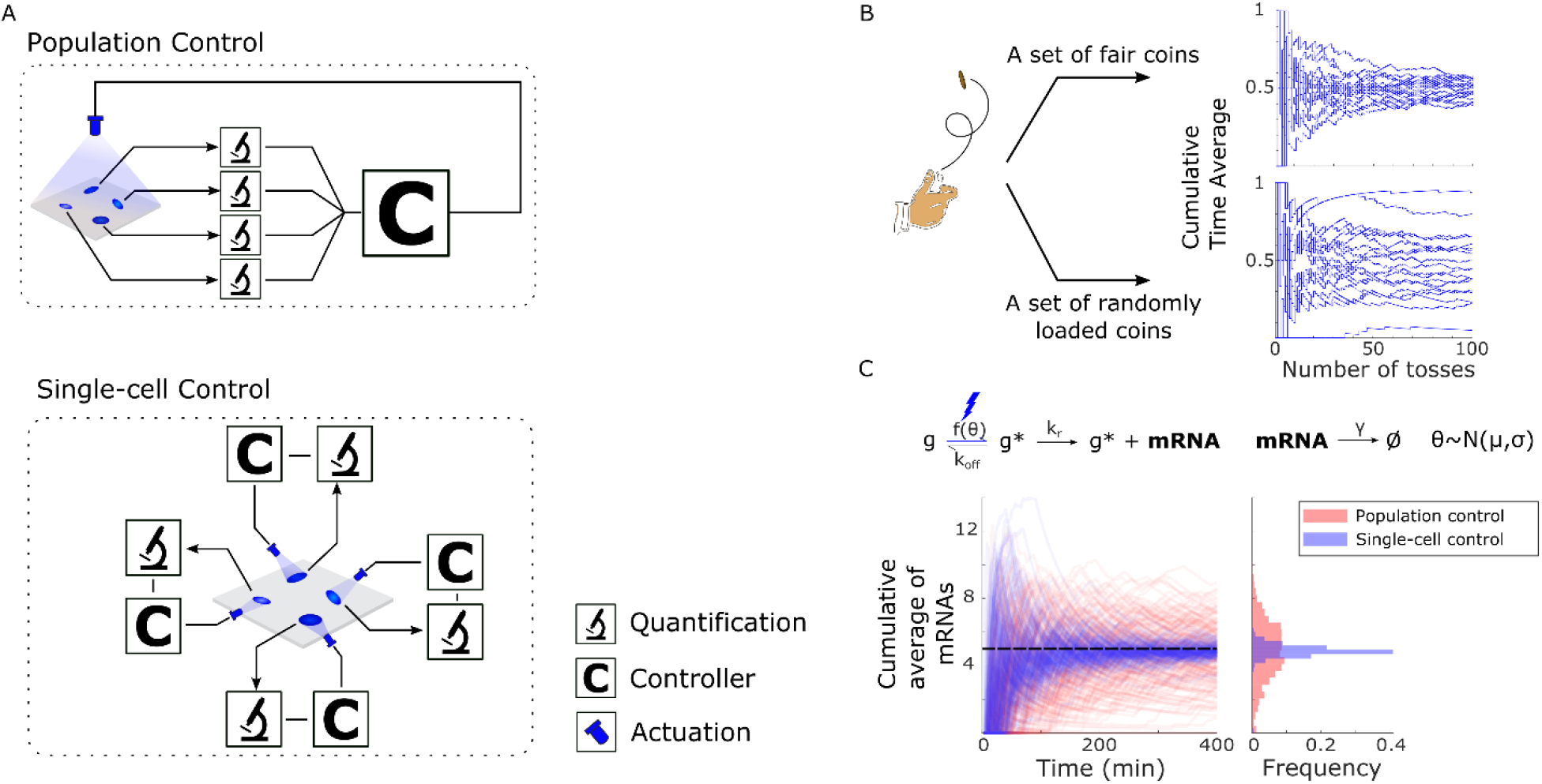
Conceptual framework: control strategies, extrinsic variation quantification. *A: Strategies for feedback control. In population control, the measurements of all cells are pooled together, generating a measure of the average cell behavior. This bulk signal is then fed to a controller, which decides on a common input to be applied to all the cells. In contrast, single-cell control generates an independent feedback loop around each cell. B: The Cumulative Time Average exposes the effect of slowly-varying extrinsic variation. To demonstrate how the CTA can be used to estimate the contribution of a static source of randomness to the total variability of stochastic system we consider a simple coin-tossing example.* As *is expected for a collection of fair coins, as the number of tosses grows, the CTAs of heads occurrences approaches 50% for all coins. Consequently, the variance of the group of CTAs progressively shrinks with the number of throws down to an asymptotic value of zero. However, if we introduce an external source of variation by assigning to each coin a randomly selected probability of landing heads, a different picture emerges. In this case, the variance of the group of CTAs reaches a non-zero steady-state value. The asymptotic value of this variance reveals information about the loading probability distribution (see section on variation decomposition). C: Reduction of extrinsic variation by single-cell control. To showcase the difference between population and single-cell control, we considered a simple stochastic model of transcription with a two-state promoter and simulated the effect of both control strategies on mRNA abundance. The transition rate from the inactive to the active promoter state is set to be a function of an external input u (e.g. light) and a parameter θ that varies from cell to cell, thereby introducing extrinsic variation into the system. By using an integral controller and simulating a collection of cells under both population (red) and single-cell (blue) feedback control, we obtain the mRNA CTAs of different cells, with the desired target value indicated by the dotted line (left plot). The plot on the right shows the distribution of mRNA CTAs across the cell sample at the end of the experiment. The tighter distribution obtained by single-cell control exemplifies the ability of this strategy to compensate for extrinsic variation.*

To compare the properties of population-level and single-cell control strategies, we implemented the same controller architecture in both settings. A simple integral control scheme (Methods) was selected to close the feedback loop at a high sampling frequency^22,25^. Besides its minimal computational complexity, this ubiquitous controller has good steady-state tracking properties and can function effectively in the presence of intrinsic variations^26^ (Methods). The performance of the two control strategies was evaluated 1) by their ability to track constant references, 2) by the feedback loop stability, understood as the absence of sustained oscillations in the controlled mRNA output, and 3) by the amount (and type) of variation present in the mRNA output.

### Quantifying contributions to cell-to-cell differences

As previous studies have demonstrated, transcription in yeast is a highly stochastic process that fluctuates on a timescale of just a few minutes^27,28^. This is indeed verified by our experiments, where transcription is observed to occur in bursts of random intensity and duration within single cells (Figure 1D, Supplementary Note 2). Considering the transcription of the modified glt1 mRNA as the system of interest and the nascent mRNA count as its measured output, the observed cell-to-cell variability in transcriptional activity at the population level can be decomposed into its intrinsic and extrinsic components^8^.

Intrinsic variation is attributed to the random events associated with individual transcription steps, such as promoter state fluctuations (partly associated with EL222 (un)binding), RNA polymerase recruitment, transcription initiation and elongation etc. On the other hand, the GLT1 gene is embedded in an environment that varies from cell to cell, for example due to differences in EL222 protein levels, amount of RNA polymerases, cell size, cell age, and cell cycle stage.

While these sources of cell-to-cell variation influence the transcription events mentioned, they are *extrinsic* to them. Moreover, within a given cell they are generally expected to fluctuate much more slowly relative to the intrinsic fluctuations of the transcription process. This characterization is in agreement with several studies of eukaryotic gene expression suggesting that extrinsic gene expression noise fluctuates over timescales roughly corresponding to the cell doubling time, while intrinsic transcriptional noise evolves over much faster timescales^29–31^.

Accordingly, extrinsic sources of variation can be assumed to be roughly constant over the timescale of transcription considered here. Henceforth, these sources of variation are represented by a vector of (random) variables *Θ* that differs from cell-to-cell^24,32^, but otherwise remains unchanged within a given cell. It follows that the total variability in the output of interest (e.g. nascent mRNA) may be separated into two parts, each originating from a different source. More precisely, the Law of Total Variance allows us to write the total variance of our fluorescent readout, *Y,* at any given time as

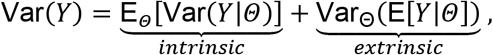

where the *Θ*-subscript denotes moments with respect to the probability distribution of extrinsic factors across the cell population, while the other moments on the right-hand side are considered with respect to all fast-varying stochastic processes within the cell.

The term labeled ‘extrinsic’ corresponds to the contribution of *Θ* to the total variance. Indeed, the conditional expectation random variable E[*Y*|*Θ*] averages out variations in *Y* due to sources other than *Θ*, and its variance with respect to *Θ* therefore captures variations in *Y* due only to variations in *Θ*. In other words, while E[*Y*|*Θ* = *θ*] measures the average readout of cells with parameter *θ,* its variance Var_Θ_(E[*Y*|*Θ*]) measures the variation of that readout due to the variation of *Θ* among cells in the population (a graphical illustration of this operation is provided in Figure 2B). On the other hand, the term labeled ‘intrinsic’ corresponds to the contribution of the fast processes to the total variance. For a given *Θ* this contribution is captured by Var(*Y*|*Θ*), and the total contributions of all cells is obtained by averaging this term over *Θ*.

With this variation decomposition in hand, one question is how the different contributions to the variance can be obtained experimentally. Var(*Y*) can be estimated from single-cell measurements of *Y* across the population. Next, we show how to estimate Var_Θ_(E[*Y*|*Θ*]) from single-cell trajectory data assuming that the intrinsic transcription process is *ergodic.* For sufficiently large times (i.e. at stationarity) the conditional expectation E[*Y*(*t*)|*Θ* = *θ*] is given by the *time average* of any sufficiently long trajectory (*y*_θ_(*t*))_*t*≥0_ of a single cell with parameter *Θ*. This is because ergodicity of the transcription process implies that such a sufficiently long trajectory contains all the information about the stationary distribution of the population of cells having the same parameter *θ.* Accordingly, we are able to approximate the average readout E[*Y*|*Θ* = *θ*] of the population of cells with parameter *θ* at stationarity by the *Cumulative Time Average* (CTA) of the trajectory (*y*_θ_(*t*))_*t*≥0_ for a sufficiently long time horizon *T*. Mathematically this CTA is defined as

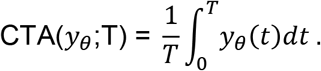

While we do not know the parameter *θ* of a given cell, the variance of these time-averages in a large ensemble of cells (corresponding to different realizations of *Θ*) gives a good estimate of Var_Θ_(E[*Y*|*Θ*]), the contribution of extrinsic factors to the total variance. Finally, by subtracting this quantity from the total population variance, the contribution of intrinsic variability can be obtained.

### Single-cell control of transcription reduces extrinsic variability

To demonstrate the capabilities of our experimental platform for optogenetic control, we used it to maintain pre-specified constant levels of transcriptional activity in yeast cells for periods of a few hours. We performed these experiments using population-level and single-cell feedback control (Supplementary Note 3). As can be observed from the CTAs of nascent mRNA counts of different cells (Figure 3A), single-cell control is capable of making all cellular readouts tightly converge around the desired values (setpoints).

**Figure 3:**
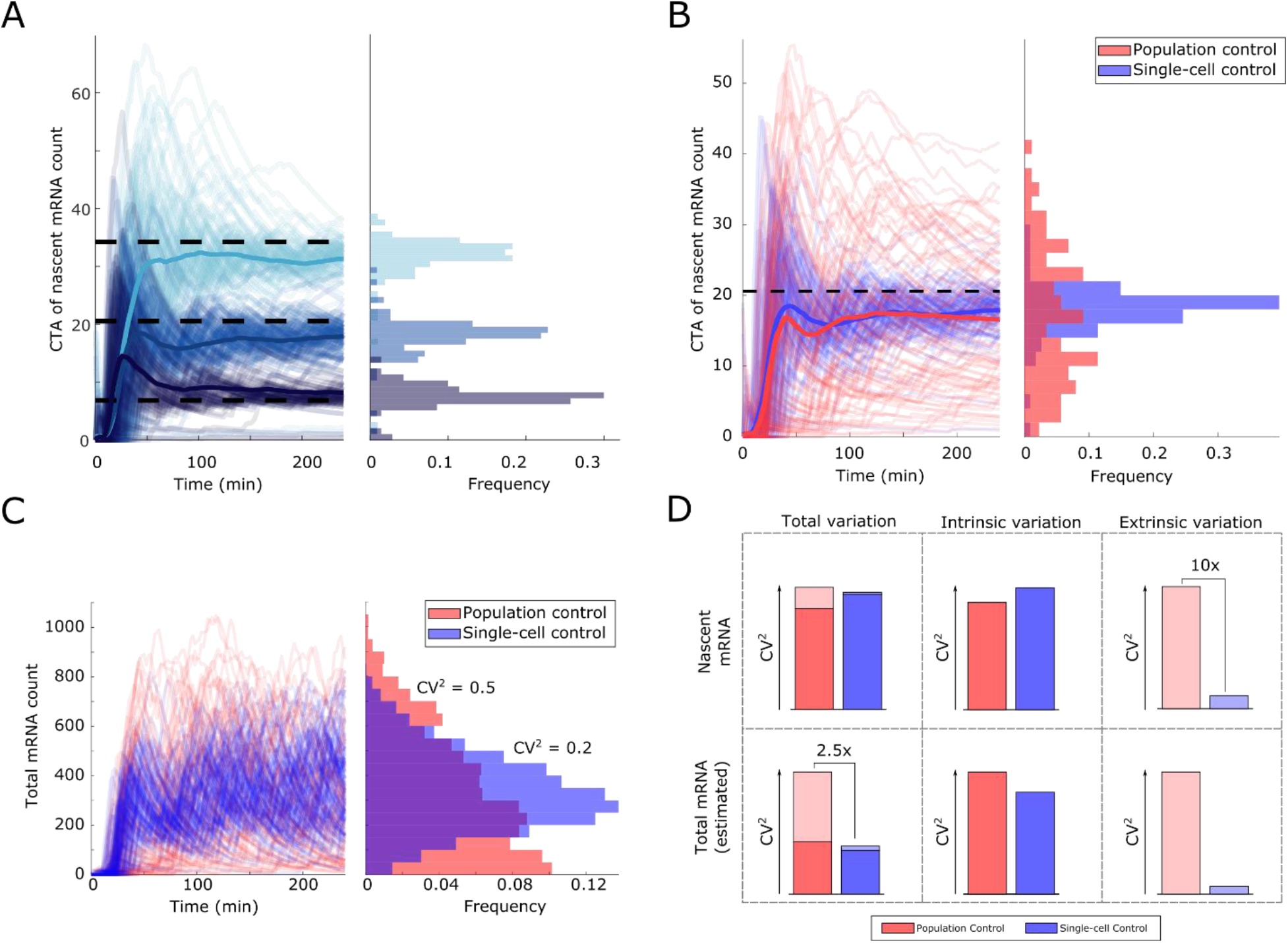
Single-cell control reduces extrinsic variation and cell-to-cell variability in mRNA numbers. *A: Tracking of constant output reference profiles. (Left) Three single-cell feedback control experiments with differing reference values (black dashed lines) show the ability of single-cell control to track constant references. Thin lines represent individual cell CTAs of nascent mRNA counts, while thick lines indicate the average behavior of the population of cells. The light input profiles can be found in Supplementary Note 3. (Right) Distribution of nascent mRNA CTAs across the cell population at the end of the experiment. B: Comparison of population (red) and single-cell control (blue). The two control strategies share the same control parameterization and reference value. (Left) Thick lines represent the average behavior of each experiment, the dashed line marks the reference value, and the thin lines are single-cell CTA traces. (Right) Distribution of nascent mRNA CTAs at the end of the experiment. The mean response (thick lines) of both experiments is nearly identical. However, the behavior of the cells in both control strategies differs considerably, with single-cell control presenting a tight distribution of the CTAs around the reference value. This result demonstrates that single-cell control is able to efficiently compensate for cell-to-cell differences, in contrast to population control. C: Estimation of mRNA counts. The number of mRNAs present in each cell was estimated using the nascent mRNA counts of the previous panel. mRNA levels obtained from single-cell control of nascent mRNA counts (blue lines) are clearly less variable than those obtained from population control (red lines). D: Variation decomposition of experimental data. The data from panel B and C is analyzed with the variation decomposition scheme described to obtain the proportion of variation attributable to extrinsic variation and intrinsic variation, for both population control (red) and single-cell control (blue). The control of nascent mRNAs in single cells reduces extrinsic variation of this variable, albeit leaving the total variation roughly unchanged with respect to population control. However, the feedback control loop in single cells clearly reduces variation in mRNA numbers. A simple model of transcription displays the same results as the experimental data.*

Comparing the CTAs of nascent mRNA counts produced by population and single-cell control (Figure 3B) reveals important differences. Even though both control strategies drive the cells to the desired setpoint on average, CTAs obtained from population control display a 10-fold greater variation (defined as the variance divided by the mean squared; CV^2^) compared to single-cell control (Figure 3D; quantification details in Supplementary Note 4). Single-cell integral feedback control achieves this dramatic reduction in extrinsic cell-to-cell variation of nascent mRNA by effectively compensating for the impact of the constant (or slowly-varying) sources of such variation. This is in fact a robustness feature of integral feedback – one that is related to its ability to achieve perfect rejection of constant disturbances in the steady-state^25^. On the other hand, the same single-cell integral feedback control does not reduce intrinsic variation of nascent mRNA and even slightly increases it in comparison to population feedback (or constant input) that gives rise to the same mean. In fact, it can be shown analytically that for the stochastic transcription model shown in Figure 2C, single-cell integral feedback cannot reduce intrinsic variation in nascent mRNA *regardless* of the choice of controller or model parameters. This was also confirmed in simulations when the exponential waiting time for nascent mRNA release was replaced by a constant time delay that models mRNA elongation time.

The total variation of nascent mRNA consists of both intrinsic and extrinsic contributions. It is interesting to observe that extrinsic variation is only a relatively small fraction of the total nascent mRNA variation in this system. In view of the pronounced bursting observed at the single-cell level (Figure 1D), this observation can be explained by the large amount of intrinsic stochasticity present in the transcription process^7,20^. Since the intrinsic contributions dominate, the total variation in nascent mRNA is roughly similar for both control strategies (Figure 3D). This outcome, as well as the performance of the two control strategies with respect to intrinsic and extrinsic variation, were reproduced by our stochastic computer model of transcription (Supplementary Note 5). In spite of the fact that the two control strategies end up with roughly the same variation in nascent mRNA, they treat the extrinsic variation in starkly different ways. This has significant implications for the cell-to-cell variability in total mRNA, as will be demonstrated next.

In most cases of biological relevance, it is expected that the variable of interest will be the total cellular mRNA, whose abundance may need to be regulated at a desired level. Given the single-cell measurements of nascent mRNA counts collected during our feedback control experiments, we can also estimate the total number of mRNAs present in each cell over time. To do this, two additional parameters are required: the average amount of time it takes for a nascent mRNA to finish transcription and diffuse away from the transcription site, and the average mRNA half-life. Both parameters can be relatively easily obtained experimentally. For this study, we assumed an average completion time of 2 minutes^20^ and a half-life of 25 minutes (based on the median half-life of yeast mRNAs^33^) to estimate computationally the total number of mRNAs in each cell over time (Methods). A look at the mRNAs variation statistics produced by the two control strategies reveals that total variation would be significantly reduced by single-cell control (Figure 3C, D).

This reduction of total variation at the mRNA level can be attributed to the relatively long half-life of these molecules. The mRNA dynamics thus act as a low-pass filter, removing high-frequency fluctuations arising from intrinsic processes. In contrast, extrinsic variation is not impacted by this filtering effect owing to its much slower rate of change. Therefore, the dramatic reduction of extrinsic variation in nascent mRNA count by single-cell control translates into lower total variation at the mRNA level in comparison to population control. The attenuation of mRNA variations is 2.5-fold for mRNA half-life of 25 minutes, but would increase further for longer mRNA half-lives. Again, our simple stochastic model of transcription reproduces this behavior (Supplementary Note 5, 6).

### Single-cell control improves controller performance

It is well-known that a feedback loop containing an integral controller will track constant reference inputs with zero steady-state error and be completely insensitive to constant perturbations^26^. On the other hand, integral controllers may introduce instability into the control loop even when the controlled system is not oscillatory by itself. Degrading the controller performance, these oscillations depend on the feedback gain, *K_I_*. This parameter determines how aggressively the controller responds to discrepancies between the reference value and the system output. As *K_I_* increases from zero, the rise time of the system output typically gets reduced, but at the same time the overshoot and settling time can increase to the point where the output does not converge and displays undamped oscillations.

We first explored the effect of varying the controller gain, *K_I_*, in simulation using our simple stochastic model of transcription (Figure 4A). We observed that increasing *K_I_* improves the rise time of the output in a similar manner for both control strategies. However, the feedback loop becomes oscillatory at high gains in the case of population control, implying that the controller needs to be carefully tuned to achieve good performance. This behavior is exacerbated when a delay is added to the system dynamics, which further shrinks the optimal gain range. In stark contrast, single-cell control shows no signs of instability at either high gains or delay times.

**Figure 4:**
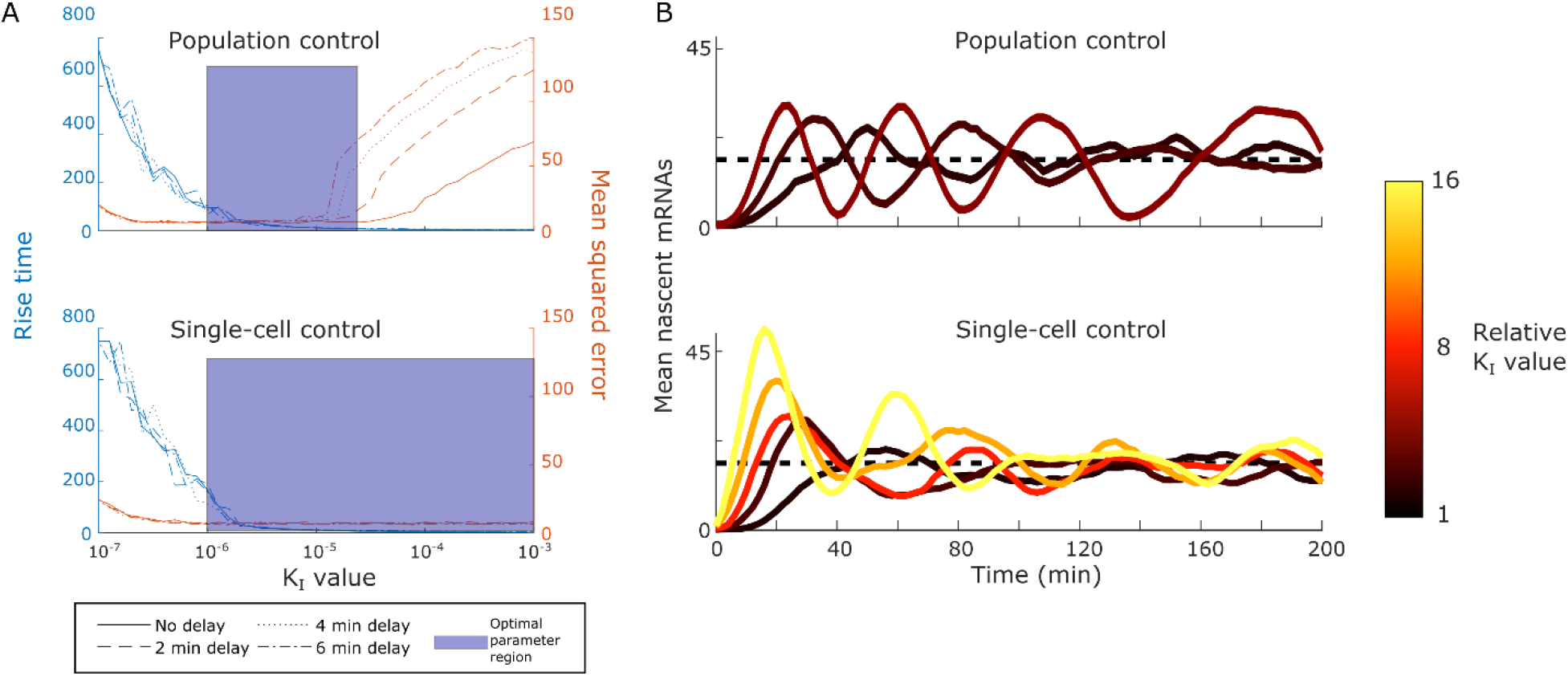
Single-cell control increases feedback loop stability. *A: In silico comparison of population control (top) and single-cell control (bottom) stability and rise time. The two control strategies are simulated under varying *K_I_* values. The stability of the feedback loop is measured by the mean squared error (orange) between the response of the system and the reference. The rise time is evaluated by the number of time-steps (blue) required to reach the reference value. Simulations of both control strategies display very similar relationship between *K_I_* value and controller responsiveness, measured by the rise time. However, population control becomes unstable at high *K_I_* values. In contrast, single-cell control does not show instability, even in presence of large delays (shown with line style) between the application of an input and the response of the system. B: Experimental results show greater stability of single-cell control. Population control causes oscillations for high *K_I_* values, while single-cell control remains stable over a wide range of controller parameterizations. Each line represents an independent experiment. The controller parameterization for each line is indicated by its color. Dark colors represent low *K_I_* values, while light colors indicate the integral controller has a high gain. The relationship between *K_I_* value and rise time is similar between both control strategies, in accordance with the simulation results.*

Experimental results of tracking constant references for varying controller parameterizations show the same trend: the population-control feedback loop becomes oscillatory after increasing the controller gain 4-fold above a baseline, while single-cell control shows convergence for *K_I_* values spanning a 16-fold difference (Figure 4B). These results are in accordance with previous theoretical analysis of similar control architectures^25^, where *in vivo* (single-cell) integral control was found to be stable in a broad range of conditions, including in stochastic settings. The larger space of well-performing controllers implies that single-cell control is less sensitive to parameter variations of the regulated system and therefore more robust.

## Discussion

We have presented an experimental platform for the independent optogenetic feedback control of more than a hundred yeast cells in parallel, consisting of a light delivery system capable of targeting light to individual cells and software to automate the measurement and tracking of S. *cerevisiae.* The platform has been tested and validated by controlling transcription, a process that displays a large degree of stochasticity in single cells. Furthermore, we introduced a new approach to separate the contributions of slowly-varying extrinsic factors and fast intrinsic fluctuations to the total variability of the system output, which we used to compare the variation-shaping properties of single-cell and population control. This analysis revealed that single-cell control is able to significantly reduce total cell-to-cell variability in mRNA numbers in comparison to population control, while offering superior dynamical performance.

Our variation decomposition approach rests on the assumption that intrinsic and extrinsic fluctuations evolve on well-separated timescales. We expect this assumption to hold in the case of transcription, which fluctuates much faster compared to changes in the cellular environment. Given this premise, our approach offers a simpler alternative to the ubiquitous dual-reporter method for variation decomposition^34,35^, in which the covariance between two identical and independent reporters is monitored. While conceptually elegant, the practical implementation of this method requires careful tuning of the two reporter systems to ensure their identical behavior and any deviations from this condition can lead to biased results.

Besides allowing the quantification of intrinsic and extrinsic variation, our experimental platform can be used to study the phenotypic consequences of gene expression variability by artificially altering the endogenous variation levels of genes of interest^36,37^. Moreover, by regulating all cellular outputs to the same target levels, our integral feedback controllers effectively “cancel out” the effect of slowly-varying sources of extrinsic variation. Analysis of the applied control inputs to each cell can thus provide information on the distribution of these “hidden variables” contributing to extrinsic variation at the population level^30,38^ and improve the predictability of cellular processes at the single-cell level.

Apart from integral feedback, our platform can easily implement alternative feedback control strategies and thus facilitate the testing of *in vivo* synthetic feedback controllers^25,39^, which are very time-consuming to implement and tune. More broadly, the setup presented here can be used to robustly drive single-cell behavior in particular regions of the output space and thus efficiently characterize gene network dynamics. It should also enable the study of systems generating toxic byproducts by regulating their concentration within pre-specified bounds to study their effects while ensuring cell viability^40^. Clamping concentrations of key regulatory molecules in single cells should also facilitate the dissection of intracellular feedback circuits^41^. Finally, one can envision the use of single-cell feedback for the spatial control of multicellular systems, such as the targeted differentiation of mammalian cells for tissue regeneration.

## Methods

### Plasmid construction

*E. coli* TOP10 cells (Invitrogen) were used for plasmid cloning and propagation. Sequences and details of all DNA constructs used in this study can be found in Supplementary Note 7. All plasmids used in this study are summarized in Supplementary Table 1. Plasmids were constructed by restriction-ligation cloning using enzymes from New England Biolabs (USA).

All PCRs were performed using Phusion Polymerase. Plasmid pDB96 is used to insert an EL222-responsive promoter and 24 PP7 stem-loops in front of the genomic GLT1 ORF and was constructed by replacing the Pol1 promoter in pDZ306^20^ with the synthetic, EL222-responsive promoter 5xELbs-CYC180 promoter (described and characterized in another manuscript, under preparation). A construct containing two copies of PCP, the SV40 NLS, and two copies of mRuby3^42^ (tdPCP-NLS-tdmRuby3) under the control of the Met25 promoter was inserted into the integrating plasmid pRG206^43^ (pDB97). All constructs were verified by Sanger sequencing (Microsynth AG, Switzerland).

### Yeast strain construction

All strains are derived from BY4741 and BY4742 (Euroscarf, Germany). All strains used in this study are summarized in Supplementary Table 2. Transformations were performed with the standard lithium acetate method^44^ and selection was performed on appropriate selection plates. DBY80, containing an EL222-responsive promoter and 24 PP7 stem-loops upstream of the GLT1 ORF, was constructed by transforming the PacI digested plasmid pDB96 into DBY41 (BY4741 expressing VP-EL222 from the Act1 promoter, construction and characterization are described in another manuscript, under preparation). A strain expressing the tdPCP-NLS-tdmRuby3 construct was generated by transforming AscI digested plasmid pDB97 into BY4742. The final strain, which is used in all experiments described in this study (DBY96), was generated by mating DBY80 and DBY91. Diploid cells were selected by growth on SD plates lacking both L-Lysine and L-Methionine.

### Culture media

Cells were grown in SD dropout medium (2% Glucose, low fluorescence yeast nitrogen base (Formedium)) with limiting concentrations of methionine (32mg/L). The medium’s pH was set to 5.8.

### Single molecule FISH experiments

For single molecule FISH (smFISH) experiments DBY96 was grown from a single colony to saturation in SD medium (32mg/L, L-Methionine). Cultures were diluted to reach an OD_700_ of 0.4 at the start of the experiment the next day. For each experimental condition, 4 ml cell culture were transferred to 25 ml glass centrifuge tubes (Schott 2160114, Duran) stirred with 3 × 8 mm magnetic stir bars (13.1120.02, Huberlab). Illumination at two different light intensities (210 and 420 μW/cm^2^, measured at 4 cm distance from the LED light source using a NOVA power meter and a PD300 photodiode sensor (Ophir Optronics)) was performed continuously using a setup comprised of a water bath (ED (v.2) THERM60, Julabo) set to 30 °C, a multi position magnetic stirrer (Telesystem 15, Thermo scientific), a 3D printed, custom-made 15-tube holder, and custom-made LED pads located underneath the culture tubes. Cultures were diluted 1:1 in fresh medium after 2 h.

Cell fixation and probe hybridization was performed as described previously^45^. Briefly, after 0, 1, 2, and 4 h of illumination, cells were fixated for 45 min after adding 400 μl of 37% formaldehyde (Sigma Aldrich) to the culture medium. Spheroplasting was performed using a final Lyticase (Sigma-Aldrich) concentration of 50 Units/ml. The progress of spheroplasting was monitored under the microscope. Cells were stored in 70% ethanol at 4 °C overnight. Hybridization was performed using multiple probes complementary to the PP7 SL and singly labeled with CY3 at a 0.1 μM concentration (synthesized by Integrated DNA Technologies, sequences are listed in Supplementary Table 3).^46^ Cells were stained with DAPI (0.1 μg/ml in PBS, Sigma-Aldrich), attached to Poly-D-Lysine treated coverslips, and slips were mounted on slides using Prolong Gold mounting medium (Invitrogen).

### Growth conditions and loading to microfluidic chip

#### Cell initialization protocol

Cell cultures were started from a -80 °C glycerol stock at least 24h prior to the experiment, and kept at OD600 < 0.2 for the last 12h leading to the experiment.

#### Microfluidic chip loading protocol

The cell culture was concentrated to an OD_600_ ~ 2 by centrifuging the sample at 3000g for 6 minutes, and discarding the appropriate volume of supernatant to reach the targeted OD600. Meanwhile, the PDMS device and cover glass (Menzel-Glaser, Germany) were rinsed with acetone, isopropanol, deionized water and dried using an air gun. The cells were then resuspended and 0.4 μl of cell solution was loaded into each chamber of the clean microfluidic chip, using a conventional pipette. The cover glass was placed on top of the PDMS device and slightly pressed down, allowing the PDMS and glass to bond electrostatically.

The loaded microfluidic chip was placed onto a custom-built microscope holder, inside the microscope’s environmental box (Life Imaging Services, Switzerland). A flow of media of at least 10 μl/min was supplied through the device via gravity flow, and the cells were allowed to settle in the new conditions for 2 hours prior to the start of any experiment.

### Image acquisition

All images were taken with a Nikon Ti-Eclipse inverted microscope (Nikon Instruments), equipped with a 40x, oil-immersion objective (MRH01401, Nikon AG, Egg, Switzerland), Spectra X Light Engine fluorescence excitation light source (Lumencor, Beaverton, USA), pE-100 bright-field light source (CoolLED Ltd., UK), and CMOS camera ORCA-Flash4.0 (Hamamatsu Photonic, Solothurn, Switzerland). The camera was water-cooled with a refrigerated bath circulator (A25 Refrigerated Circulator, Thermo Scientific). The temperature was regulated to 30 °C by an opaque environmental box (Life Imaging Services, Switzerland), which also shielded the cell sample from external light. The microscope was operated by the open-source software YouScope^47^.

All measurements were run with a diffusor and a green interference filter placed in the bright-field light path. The perfect focus system of the microscope was enabled for all measurements.

#### Fluorescence imaging

Excitation of mRuby3 was performed by the 550/15nm line from the fluorescence light source. The filter-cube used had excitation filter 561/4nm, beam splitter HC-BS573, and emission filter 605/40nm, all from AHF Analysetechnik AG (Tubingen, Germany). Image stacks with approximately 0.5 μm z steps were taken with an exposure time of 300ms per image. With these imaging settings, images can be taken every 2 minutes for a period up to 4h without bleaching more than 15% of the initial cell fluorescence (Supplementary Note 8).

#### Microscopy setting for smFISH

smFISH images were acquired using a Plan Apo Lambda 100X Oil objective (Nikon Instruments). Z-stacks consisting of 31 images with a step size of 0.1 μm were taken for CY3 (Excitation: 542/33, Emission: 595/50) and DAPI (Excitation: 390/22, Emission: 460/50). Phase contrast images were taken at the reference point of the Z-stacks to allow for cell segmentation.

### Image analysis

#### Cell segmentation and tracking

Bright-field images below and above the focal plane (Nikon Perfect Focus System, +/-5 AU) were acquired for cell segmentation and tracking. The image above the focal plane was divided by the one below the focal plane to eliminate uneven illumination and enhance the border of the cells. Segmentation was performed on the resulting image using Matlab (MathWorks) code extracted from the CellX software tool^48^. Cell tracking from frame to frame was accomplished with Matlab scripts based on ref. 49.

#### Quantification of nascent mRNAs

In our experimental setup, nascent mRNAs can be visualized in the Cy3 fluorescence channel, and appear as a diffraction-limited spot, as they accumulate at the transcription start site. The fluorescence intensity of a diffraction-limited spot can be described by an Airy pattern, whose central lobe is well approximated by a Gaussian function. Under this approximation, the volume of the Gaussian function is proportional to the number of nascent mRNAs constituting the fluorescent spot.

To quantify the number of nascent mRNAs in each cell we take a z-stack of fluorescent images, spanning the whole cell volume. For each captured image, we perform the following analysis:

1. We first remove the fluorescent background signal by means of a Gaussian filter. The Gaussian filter clears features smaller than its standard deviation
2. Next we subtract the original image by the filtered image, obtaining a third image where the fluorescent background has been removed, while preserving features of the size of the fluorescent spots we wish to quantify
3. Finally, we fit a 2D Gaussian function to the pixel intensity surface of each cell (Supplementary Note 9). Two measures of the goodness of fit of the fitted Gaussian function, as well as its standard deviation and amplitude are used to classify a cell as either being transcriptionally-active or inactive (Supplementary Note 9).

Multiple diffraction-limited spots can be detected in one cell, because of the signal overlap between consecutive fluorescent images in the z-stack. If this happens, the spot with the highest signal is taken as the measurement of nascent mRNA for that cell.

### Calibration of spot intensities to number of nascent mRNAs

The conversion factor between fluorescent spot intensities (a.u.) and number of nascent mRNAs was measured in accordance to ref. 7. The distribution of spot intensities obtained for a constant light intensity of the DMD projector was aligned to the distribution of nascent mRNAs as quantified by smFISH, performed on the same yeast strain and also exposed to constant light conditions. The percentiles of each distribution were used as calibration points for the alignment. More details are present in Supplementary Note 10.

### Light-delivery system

#### Hardware

Optogenetic stimulation was done with a DMD projector (DLP LightCrafter 4500, Texas Instruments) mounted on an optical table, together with the necessary optical elements to focus the emitted light at the focal plane of the microscope’s objective. A schematic of the setup, together with a list of components is provided in Supplementary Note 1 and Supplementary Table 4, respectively. The light intensity at the specimen and the blue-light spectra is shown in Supplementary Note 1.

#### Projection image correction

The light-delivery system was aligned to the microscope’s camera prior to the start of each experiment. This procedure consists of finding the correspondence between projector pixels and camera pixels. The knowledge of this mapping is required to precisely target with light the cells in the field-of-view. The calibration procedure is described in Supplementary Note 1.

### Fabrication of microfluidic device

The microfluidic chip, adapted from ref. 50 was fabricated as described. The chip is a single layer poly(dimethylsiloxane) (PDMS, Sylgard 184, Dow Corning Corp., USA) device, attached to a cover glass (thickness: 150 μm, size: 24 mm x 60 mm).

### Modelling

#### Gene expression model

We model transcription using a two-state promoter model^51–53^. The reaction rate for the transition of the promoter conformation from closed (G) to open (G*) has been replaced by a function dependent on the active amount of transcription factor (TF*). The fraction of active transcription factor in turn depends on the light input given to the system (u).

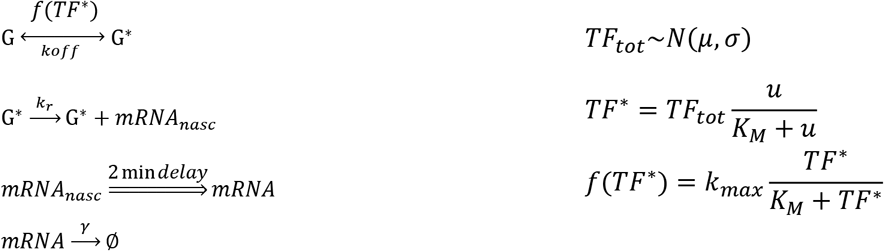

Nascent mRNAs are only produced from the open promoter conformation (G*). Given the relatively tight distribution of dwell times of individual nascent RNAs, we assume that the dwell time of each nascent RNA at the transcription site is equal to two minutes^20^. In this way, in our model a nascent mRNA is converted to a mature mRNA two minutes after its birth event.

Extrinsic variation is introduced into the model by assigning different total amounts of transcription factor, TFtot, to each cell. The parameter is drawn from a Gaussian distribution truncated at 0, and is set to remain constant for the duration of the experiment. As the external input (u) determines the fraction of active transcription factor (TF*), cells with more TF_tot_ will present a stronger response to a given input.

All parameters used in the simulation are found in Supplementary Table 5. For the closed-loop simulations, the light input (u) was updated every 2 minutes (the same frequency used in the feedback control experiments). For population control, the readouts of all simulated cells were averaged and fed into a common controller. Delay was added to the closed-loop simulations by delaying the application of the input for the appropriate number of time-steps.

#### Estimation of mRNA abundance from nascent mRNA time-course data

The mRNA abundance for each cell was estimated from the experimentally measured nascent mRNAs time-course, based on a simple mathematical model. According to this, the conversion of each newly generated nascent RNA to mature mRNA is assumed to take place two minutes after the appearance of the nascent molecule. Furthermore, the life time of each created mRNA follows an exponential distribution. This mean that the probability of an mRNA degradation event between two measurements is

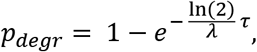

where *λ* is the mRNA half-life, and *τ* is the sampling period.

#### Stochastic simulations

All simulations were performed with Matlab (Mathworks), using the Random Time Change (RTC) algorithm^54^.

### Description of control algorithms

To regulate the number of nascent RNAs to a desired constant reference value, we used integral feedback controllers^26^ both for single-cell and population control. In integral control, the input applied to the controlled system is proportional to the integral of the output error. In our experiments, the controller output (applied blue light intensity) is updated once a new output measurement becomes available, and is held constant between measurement times.

More specifically, given the system output at measurement time *t_k_*, *y*(*t_k_*), and the desired output reference value *y_ref_*, the error *e*(*t_k_*) = *y_ref_ – y*(*t_k_*) is formed and the controller output, *I*(*t_k_*), is defined as 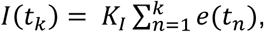, where *K_I_* denotes the *controller gain.* By adjusting this parameter, the controller can be tuned to respond more or less aggressively to output deviations from the desired reference. In our experiments, the controller gain was chosen through manual tuning and kept the same for the two control strategies.

Due to the fact that negative inputs have no physical meaning and the projector output has an upper power limit, the applied input to the system at time *t_k_, u(t_k_),* is given by

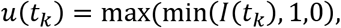

where 1 corresponds to the maximum (scaled) light intensity that the projector can provide.

In the case of single-cell control, *y*(*t_k_*) and *u*(*t_k_*) correspond to the output and applied input of a single cell respectively, since each cell is controlled by separate integral controller. For population control, the individual cell outputs over the cell population are pooled together and averaged. The computed mean is then fed to a single integral controller which computes one common input for all cells.

It is a well-known fact in automatic control theory that in a stable *deterministic* feedback loop containing an integral controller, the steady-state system output will be equal to the reference value^26^. This can be easily seen by the fact that *I(t)* will stop changing only when the error converges to zero. This analysis is applicable in the case of population control, where the population mean is the controlled output and follows deterministic dynamics.

When the controlled system is *stochastic* (as in the case of single-cell feedback), provided the closed-loop system converges to a unique stationary distribution (the equivalent of a unique stable equilibrium point for deterministic systems) then the output *mean* should again be equal to the reference. In the opposite case, the average error would be non-zero and the controller output would not be stationary.

*Note: all design schematics, computer code and experimental data are available from the authors upon request.*

## Acknowledgements

The authors would like to thank Andreas Hierlemann (ETH Zurich) for providing access to the cleanroom facilities, and Fabian Rudolf (ETH Zurich) for the cell tracking code and plasmids. We would also like to thank Michael Lin (Stanford University), Robert H Singer (Albert Einstein College of Medicine), Daniel Zenklusen (Université de Montréal), and Kevin Gardner (City University of New York) for providing plasmids.

## Author Contributions

M.K. and A.M.-A conceived the project. M.R. developed the automation and control software, built the experimental platform, performed the feedback control experiments. M.K. A.M.-A, M.R. and A.G. analyzed the data. D.B. designed and built plasmids and strains, provided technical assistance, and performed all smFISH assays. G.W.S. designed and produced the microfluidic chip, and provided helpful advice on the microscopy imaging setup. A.G. formulated the variation decomposition. M.K. and A.M.-A supervised the project. M.K., A.M.-A, and M.R wrote the manuscript, with valuable input from A.G. and D.B.

